# Root type and soil phosphate determine the taxonomic landscape of colonizing fungi and the transcriptome of field-grown maize roots

**DOI:** 10.1101/198283

**Authors:** Peng Yu, Chao Wang, Jutta A. Baldauf, Huanhuan Tai, Caroline Gutjahr, Frank Hochholdinger, Chunjian Li

## Abstract

Our data illustrates for the first time that root type identity and phosphate availability determine the community composition of colonizing fungi and shape the transcriptomic response of the maize root system.

**Summary:**

- Plant root systems consist of different root types colonized by a myriad of soil microorganisms including fungi, which influence plant health and performance. The distinct functional and metabolic characteristics of these root types may influence root type inhabiting fungal communities.
- We performed internal transcribed spacer (ITS) DNA profiling to determine the composition of fungal communities in field-grown axial and lateral roots of maize (*Zea mays* L.) and in response to two different soil phosphate (P) regimes. In parallel, these root types were subjected to transcriptome profiling by RNA-Seq.
- We demonstrated that fungal communities were influenced by soil P levels in a root type-specific manner. Moreover, maize transcriptome sequencing revealed root type-specific shifts in cell wall metabolism and defense gene expression in response to high phosphate. Furthermore, lateral roots specifically accumulated defense related transcripts at high P levels. This observation was correlated with a shift in fungal community composition including a reduction of colonization by arbuscular mycorrhiza fungi as observed in ITS sequence data and microscopic evaluation of root colonization.
- Our findings point towards a diversity of functional niches within root systems, which dynamically change in response to soil nutrients. Our study provides new insights for understanding root-microbiota interactions of individual root types to environmental stimuli aiming to improve plant growth and fitness.

## Introduction

Land plants host a wide variety of root-inhabiting microbes (Bulgarelli *et al*., 2013). These microorganisms substantially support their host plants in the acquisition of soil nutrients (Smith & Smith, 2011). Moreover, the microbiota contributes to plant health by suppressing pathogens or enhancing disease resistance (Mendes *et al*., 2011; Berendsen *et al*., 2012). Studies in Arabidopsis, rice and maize have shown that the taxonomic composition of the root inhabiting microbiota are strongly influenced by geography and soil types (Bulgarelli *et al*., 2012; Lundberg *et al*., 2012; Edwards *et al*., 2015), but also by the plant genotype (Aira *et al*., 2010; Bouffaud *et al*., 2012; Bulgarelli *et al*. 2012; Edwards *et al*., 2015). Thus, it is possible that plants attract microbes, which are most beneficial to them under a given environmental condition.

Maize is one of the most important cereal crops (Gore *et al*., 2009). Its complex root architecture and morphology is substantially influenced by environmental variation of soil conditions (Yu *et al*., 2016a). In cereal crops, highly branched root systems are composed of multiple root types formed at different developmental stages under the control of distinct genes (Tai *et al*., 2016). Axial roots generally function in conferring anchorage to the soil while the finer, soil-exploring lateral roots are mainly involved in foraging nutrients and water resources (Coudert *et al*., 2010; Rogers & Benfey, 2015; Yu *et al*., 2016a). The observed “job sharing” among root types within a root system implies root type-specific molecular and physiological responses to biotic and abiotic stimuli. In fact, recent root type-specific transcriptomic and lateral root branching responses to local high nitrate have been described in maize (Yu *et al*., 2015, 2016b). Moreover, the colonization of root systems by soil microbes such as arbuscular mycorrhiza fungi (AMF) is uneven among root types. This has been well described in rice, in which large lateral roots are strongly colonized, whereas crown roots are only slightly colonized and fine lateral roots are not colonized (Gutjahr *et al*., 2009; Gutjahr & Paszkowski, 2013; Fiorilli *et al*. 2015). These differences are mirrored by distinct transcriptional profiles among rice root types during AM symbiosis, suggesting potential relationships between root colonization, architectural variations and functional switches within the root system (Gutjahr *et al*., 2015).

Phosphorus (P) is one of the most limiting resources in natural soils and its availability is critical for crop productivity. Plant roots can absorb only inorganic orthophosphate (Pi), although P is abundant in many natural soil types both as organic and inorganic pools (Marschner *et al*., 2011). Plants have evolved molecular systems that can sense and respond to P starvation and adjust root and shoot growth accordingly (Poirier & Bucher, 2002). In addition, as a mechanism to adjust internal phosphate homeostasis, roots are associated with bacteria and fungi, which can mobilize inorganic P in soils inaccessible for plants such as hydroxyapatite and Ca_3_(PO_4_)_2_ by conversion into bioavailable P (Smith & Read, 2008). AMF of the phylum Glomeromycota are the best-characterized beneficial fungi associated with plant roots. They colonize 80%-90% of terrestrial plants and take up soil nutrients, including poorly mobile P, via an extended extraradical hyphal network (Bonfante & Genre, 2010; Smith & Smith, 2011; Gutjahr & Paszkowski, 2013). These nutrients are then transported into the root and released at highly branched hyphal structures, the arbuscules, which form inside root cortical cells (Smith & Smith, 2011; Gutjahr & Parniske, 2013). Besides AMF, roots are also inhabited by fungi from other phyla such as Ascomycota, Zygomycota and Basidiomycota. Some members of these fungal clades form ectomycorrhizas with woody plants (Vrålstad, 2004; Smith *et al*., 2007) and others live in roots or in the rhizosphere as endophytes or pathogens (Ambardar *et al*., 2016). Less is known about non-AMF fungal communities in roots and rhizospheres, although several studies cataloging the root and rhizosphere bacterial microbiome or AMF communities have been conducted.

High-throughput sequencing technologies have facilitated systemic surveys of root-associated microbiomes and interactions with their habitats (Ofek-Lalzar *et al*., 2014). Recently root type-specific regulation of root system architecture has been surveyed at the transcriptional level (Gutjahr *et al*., 2015; Yu *et al*., 2016b). However, interaction between transcriptome changes and the interior fungal community within the root types remains poorly described, especially under realistic field conditions. In this study, taxonomic identification of fungal communities inhabiting different root types by ITS DNA amplicon sequencing was combined with transcriptome analyses of these root types by RNA sequencing (RNA-Seq). The results highlight root type-specific fungal taxonomic compositions and transcriptome profiles in response to divergent P regimes. Our findings point towards a diversity of niches within the root system, which dynamically change in response to environmental factors such as soil nutrients.

## Materials and Methods

### Experimental design and sample collections in the field

Hybrid maize of the genotype ZD958 (a modern variety widely used in China) was planted in four biological replicate plots in the field under high (150 kg ha^-1^) and low (0 kg ha^-1^) phosphate conditions at the long-term experimental station of China Agricultural University (40°8'20"N, 116°10'047"E) in 2015. The design of this long term experiment is as follows: in total eight phosphate levels are tested and each phosphate level is represented by four independent blocks. The 32 blocks are randomly distributed across the field. In our experiment, we selected one low and one high phosphate condition. We collected each of the four biological replicates of root and soil samples for the two soil phosphate levels from a different block. The different blocks per treatment are spatially separated from each other by blocks, which were subjected to different phosphate treatments. This long term experiment and the block design is described in detail in Deng *et al*. (2014) and Wang *et al*. (2017). The soil type at the study site is a calcareous alluvial soil with a silt loam texture (FAO) typical of the north region of China. The top 30 cm soil from each independent block were pooled and mixed to determine soil chemical properties prior to sowing. The chemical properties of these soils are listed in Table S1. The maize seeds were sown on 6/5/2015 and samples for subsequent analyses were collected on 7/25/2015. The exact amounts of essential chemical fertilizers for the two phosphate treatments (low phosphate and high phosphate) were weighed and applied separately at each application date and are provided in Table S2. The average monthly rainfall across the whole field was recorded until sample collections by a small meteorological station located in the experimental field listed in Table S3.

For fungal community analyses under each phosphate condition samples of bulk soil, axial roots and lateral roots were taken at the flowering stage (Fig. 1a). After shoot excision, all mature axial roots with emerged lateral roots of two neighboring plants from the top 30 cm soil for each independent block were vigorously washed with sterilized deionized water in order to remove all soil from the root surface. The washing steps were repeated twice to avoid soil contamination in the root type samples. Subsequently the root system was separated into axial and lateral root samples with sterilized scissors. Axial roots without lateral roots and newly emerged axial roots were excluded from our study to minimize developmental variability within the pool of roots (Gutjahr *et al*., 2015). Separated axial and lateral root samples were gently dried with clean soft tissue and immediately frozen in liquid nitrogen and stored at -80 °C for downstream microbiome and transcriptome analysis. The top 30 cm soil layer at the unplanted plots was crushed and sieved through a 2 mm mesh in the field. This mixed fresh soil was referred to as the bulk soil samples and stored at -20 °C for subsequent short amplicon sequencing analyses. Shoot biomass and P content of maize plants with low or high P input were determined using a modified Kjeldahl digestion and vanado-molybdate method by automated colorimetry according to Peng *et al*. 2012 (Table S4).

**Fig. 1.**
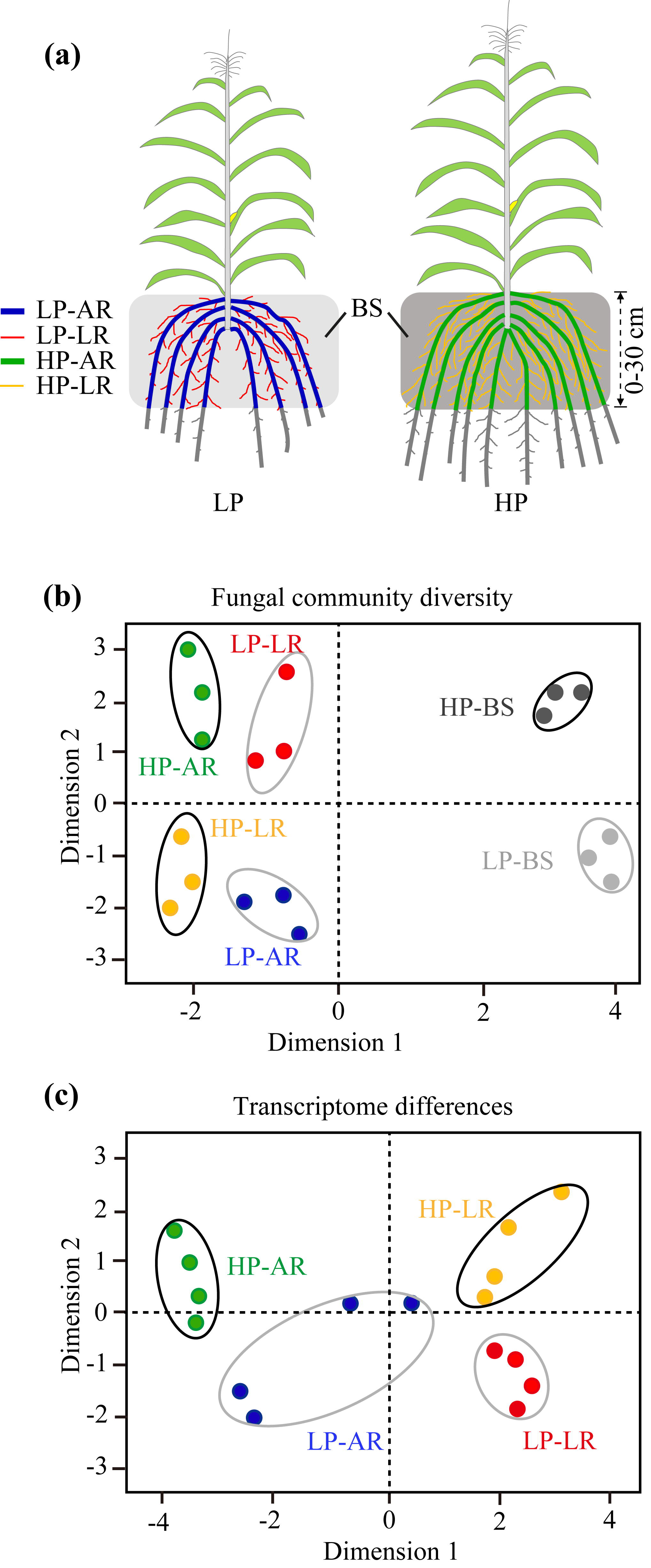
Fungal community composition and root transcriptomes differ among maize root types and soil phosphate levels. (a) Illustration of maize plants depicting the axial and lateral roots sampled from the top 30 cm soil layer. (b) Multidimensional scaling plot of fungal communities in root types and bulk soils under HP and LP as profiled by ITS gene sequencing. (c) Multidimensional scaling plot of RNA populations in maize root types under HP and LP. AR, axial roots; BS, bulk soil; HP, high phosphate; LP, low phosphate; LR, lateral roots.

### Short amplicon sequencing experiments for root types and bulk soils

#### Extraction of genomic DNA

Total genomic DNA was extracted from ground root tissues of the two root types and from bulk soil with the FastDNA® SPIN Kit (MP Biomedicals, Solon, USA) following the manufacturer’s instructions. DNA concentration and purity was estimated on 1% agarose gels. Subsequently, DNA was diluted to 1ng/µl with sterile water.

#### Amplicon generation

Fungal diversity was characterized by sequencing ITS sequences amplified by PCR from bulk soil, axial root and lateral root DNA extracts. Briefly, fungal ITS1 loci were amplified with primers ITS5-1737F (5'-GGAAGTAAAAGTCGTAACAAGG-3') and ITS2-2043R (5'-GCTGCGTTCTTCATCGATGC-3'), which are universal DNA barcode marker for fungi (Schoch *et al*., 2012) and widely used for species identification for soil fungal community (Tedersoo *et al*., 2010; Lu *et al*., 2013). These specific primers (New England Biolabs) included a barcode and adaptor for annealing to the Illumina flow cell. All pooled triplicate PCR reactions were carried out in a 30 µl volume with 15 µl of Phusion® High-Fidelity PCR Master Mix (New England Biolabs), 0.2 µmol of forward and reverse primers, and about 10 ng template DNA. Thermal cycling included an initial denaturation step at 98 °C for 1 min, followed by 30 cycles of denaturation at 98 °C for 10 s, annealing at 50 °C for 30 s, and elongation at 72 °C for 30 s and a final step of 72°C for 5 min.

#### PCR product quantification, quality control and purification

PCR products were separated on a 2% agarose gel and visualized with SYBR green. Eighteen DNA samples with bands of a size between 400-450 bp were chosen for further experiments. PCR products of eighteen samples were mixed in equidense ratios. Mixed PCR products were purified with the GeneJET Gel Extraction Kit (Thermo Scientific).

#### Library preparation and sequencing

Sequencing libraries were generated using NEB Next Ultra DNA Library Prep Kit from Illumina (NEB, USA) following the manufacturer’s instructions including the addition of index codes. Library quality was assessed on the Qubit 2.0 Fluorometer (Thermo Scientific) and High-Sensitivity DNA chip (Agilent Bioanalyzer). Finally, the library was sequenced on an Illumina HiSeq 2000 platform and 250 bp paired-end reads were generated.

### Data analysis of short amplicon sequencing

Raw Illumina fastq files were demultiplexed, quality filtered and analyzed using a custom Perl script by QIIME v1.7.0 (Caporaso *et al*., 2010; Dataset S1). Paired-end reads of the original DNA fragments were merged into raw tags by using FLASH (Magoč & Salzberg, 2011) and then assigned to each sample according to the unique barcodes. Quality filtering on the raw tags was performed under specific filtering conditions to obtain the high-quality clean tags using QIIME v1.7.0 (Bokulich *et al*., 2013). In-house Perl scripts were used to analyze alpha-(within samples) and beta-(among samples) diversity. First, reads were filtered by QIIME quality filters. Then pick_de_novo_otus.py was used to pick OTUs to generate an OTU table. Sequences with ≥97% similarity were assigned to the same OTUs using UCLUST. We picked a representative sequence for each OTU and used the Unite Database (Kõljalg *et al*., 2013) to annotate taxonomic information base on Blast algorithm which was calculated by QIIME software. OTU relative abundances were calculated by dividing the absolute abundances by the total sequence count per sample analyzed. In order to compute alpha diversity, we rarified the OTU table and calculated three metrics: Chao1 is estimated as the species abundance. Observed species are estimated as the amount of unique OTUs and the Shannon index is estimated as the diversity found in each sample. Rarefaction curve was generated based on the chao1 metric. Multidimensional scaling was performed to visualize the sample relations based on the Bray-Curtis similarity matrix using the plotMDS function of the Bioconductor package limma (Smyth, 2005) in R.

Beta diversity was calculated based on unweighted UniFrac distance by QIIME software. Unweighted pair group method with arithmetic mean (UPGMA) clustering was performed as a type of hierarchical clustering method to interpret the unweighted UniFrac distance matrix using average linkage by QIIME. Tukey’s post-hoch tests and Student’s *t*-tests were conducted to determine the fungal diversity of the different root types at a given phosphate condition. Relative abundance of fungal taxa among root type and soil samples were determined and statistical analyses were based on FDR-corrected Kruskal-Wallis test (*P* <0.05).

### Assays for arbuscular mycorrhizal fungi colonization of root types

Representative axial and 1^st^ and 2^nd^ order lateral roots selected from the root samples collected at 0-30 cm soil depth at flowering stage were stained with nonvital Trypan Blue (Shanghai Urchem Ltd, China) according to Phillips & Hayman (1970). Stained roots were studied under a microscope and the intensity of root cortex colonization by AMF was determined as described by Trouvelot *et al*. (1986). The arbuscular mycorrhizal colonization frequency represents the occurrence intensity of the sum of all AMF structures in the root samples. Arbuscule abundance denotes the arbuscule density within colonized root areas.

### Transcriptome profiling of maize root types

#### Extraction of total RNA, cDNA library construction and RNA-Seq

For each of the two root types and two P conditions frozen samples of the axial and lateral roots were ground in liquid nitrogen in four biological replicates resulting in a total of sixteen samples. RNA was extracted by the RNeasy Plus Universal Mini Kit (Qiagen). RNA quality was assessed by agarose gel electrophoresis and by an Agilent RNA 6000 Nano LabChip on an Agilent 2010 Bioanalyzer (Agilent Technologies). All RNA samples had an excellent quality as documented by RIN values from 7.6 to 8.9 (Fig. S1). During the quality control steps, an Agilent DNA 1000 LabChip (Agilent Technologies) and an ABI StepOne Plus Real-Time PCR System (Applied Biosystems) were used for quantification and quality control of the sample libraries. The cDNA libraries for RNA-Seq were constructed with the TruSeq RNA Sample Prep Kit (Illumina). For sequencing, sixteen libraries for two root types under two P levels were evenly distributed into two lanes of a flow cell. Cluster preparation and paired-end read sequencing were performed according to the HiSeq 2000 guidelines (Illumina).

#### Data processing and analysis

Processing, trimming, mapping of raw RNA-Seq reads were performed by CLC Genomics Workbench as previously described (Yu *et al*., 2015; Dataset S2). Only genes that were represented by a minimum of five mapped reads in all four replicates of at least one root type were declared expressed and considered for downstream analyses. The raw sequencing reads were normalized by sequencing depth and were log_2_-transformed to meet the assumptions of a linear model. Multidimensional scaling analysis and statistical procedures for analyzing differentially expressed genes between axial and lateral roots in combination with two P conditions were performed using the Bioconductor package *limma* (Smyth, 2004, 2005) in R (R version 3.1.1 2014-07-10, limma_3.20.9) according to Yu *et al*., 2016b. The resulting *P*-values of the pairwise *t*-tests were used to determine the total number of differentially expressed genes for each comparison by controlling the FDR <0.05 to adjust for multiple testing (Benjamini & Hochberg, 1995).

### Functional annotation and associated network analysis

MapMan software (Thimm *et al*., 2004) was used to assign and subsequently visualize differentially expressed genes to metabolic pathways based on the functional annotation file ZmB73_5b_FGS_cds_2012. Chi-Square and Fisher’s exact tests were used to determine if the observed number of genes in each of the 35 major MapMan categories significantly deviated from the expected distribution of all expressed genes (Marcon *et al*., 2015). Association networks of the identified genes significantly enriched in MapMan categories were constructed with the aid of the online analysis tool STRING v10 (Szklarczyk *et al*., 2015) and functional connections between each pair of interconnected genes were determined at a high confidence of >0.7 (Yu *et al*., 2016b). Statistical analyses of functional enrichments in the network were further determined by KEGG (Kyoto Encyclopedia of Genes and Genomes) pathways (Kanehisa *et al*., 2011) to identify significant biological processes.

## Data availability

Raw plant RNA-Seq data and fungal ITS sequencing data were deposited at the Sequence Read Archive (http://www.ncbi.nlm.nih.gov/sra) under accession numbers SRP095256 and SRP092319, respectively.

## Results

### Global patterns of root type-specific fungal communities and transcriptomes under diverse P conditions

The root system of maize consists of a variety of axial roots including primary, seminal and shoot-borne roots. All of these root-types form lateral roots. We examined the differences in fungal community composition between axial and lateral roots grown in the field under low P (LP) and high P (HP) conditions and correlated the fungal community composition with the corresponding maize root transcriptomes. Moreover, bulk soils with the two P levels were collected (Fig. 1a) for determination of free living soil fungi. Overall, fungal taxonomic structure varied across root types and P conditions, but replicate samples clustered closely together (Fig. 1b, Dataset S1). Moreover, the fungal taxonomic composition of bulk soils varied strongly between the two P levels and was very different from the fungal taxonomic composition associated with specific root types (Figs. 1b, S2).

In parallel, the transcriptomes of the axial and lateral roots were determined by RNASeq to survey gene expression in the two root types under two P levels (Datasets S2, 3). In total, 27,375 genes were expressed in at least one root type/P treatment variant (Dataset S3). A multi-dimensional scaling plot showed the distances between transcript populations of root types and P levels (Fig. 1c). It highlighted that replicate root type by P regime samples clustered together and that transcriptomic differences were more divergent among root types than among P treatments.

### Fungal taxonomic composition differs among maize root types under diverse P conditions

We identified in total 587 OTUs (operational taxonomic units), defined as general units of microbial taxonomic classifications under LP and 458 OTUs under HP conditions in bulk soil, axial and lateral root samples (Fig. 2a). The relative abundance of enriched fungal OTUs and taxonomic information are listed in Dataset S4. Among those OTUs, under LP conditions 85 were specifically detected in lateral roots and 67 in axial roots, indicating a distinct fungal taxonomic structure between the root types. Under HP conditions 53 OTUs were specific to lateral roots and 111 to axial roots. Under both P conditions, 67% of these OTUs were also found free-living in soil. However, both axial and lateral roots showed specific OTUs, which were not detected in bulk soil. Among those, 40 were exclusively enriched in lateral roots and 5 in axial roots under LP conditions. Under HP conditions 19 OTUs were exclusively enriched in lateral roots and 55 were restricted to axial roots (Fig. 2a). This indicates that the taxonomic complexity of the fungal community is co-influenced by root type, P availability and their interaction.

**Fig. 2.**
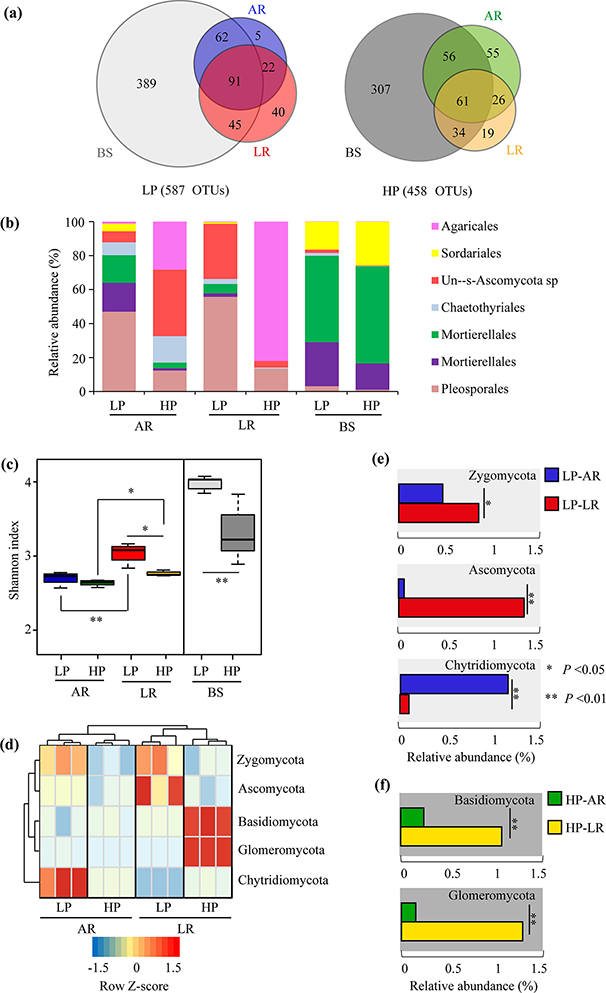
Abundance of fungal taxonomic units in maize roots and bulk soil with low-or high-P supply. (a) Numbers of differentially enriched fungal OTUs in maize roots and bulk soil. (b) Relative abundance in root types of fungal taxa at the order level. The figure shows the same distribution of significantly enriched fungal taxa as in Fig. S3, without the taxa that were highly enriched in soil and not significantly different among root types. The fungal order Mortierellales with different OTUs indicates different taxonomic identifications. Statistical significance was tested using the Kruskal-Wallis test (FDR-corrected *P* <0.05) at the order level. (c) Shannon index of fungal communities in root type and bulk soil samples at two different phosphate levels. Asterisks denote significant differences between low and high P levels for a given root type according to Tukey’s post-hoc test (\**P* <0.05; \*\**P* <0.01). Asterisks denote significant fungal diversity between two soil types according to Student’s *t* tests (\**P* <0.05; \*\**P* <0.01). (d) Hierarchical clustering analysis based on the OTUs of axial and lateral roots under LP or HP at the phylum level. Relative abundance of differentially abundant phyla are shown and normalized by *Z*-score across all datasets. The dendrogram was inferred by applying the unweighted pair group method with the arithmetic mean (UPGMA) as distance function. Distinct phyla enriched in axial and lateral roots under low (e) and high (f) P supply. AR, axial root; HP, high phosphate; LP, low phosphate; LR, lateral root; OTU, operational taxonomic unit. Asterisks denote significantly enriched phyla between axial and lateral roots according to paired Student’s *t* tests (\**P* <0.05; \*\**P* <0.01).

To understand how root type and P level influence the taxonomic structure of root inhabiting fungi, the OTUs were classified at the phylum level (Ascomycota, Basidiomycota, Glomeromycota, Zygomycota, Chytridiomycota). Because significant proportions of the microbial diversity were shared among root types, we focused on the differences in the relative abundance of taxa among root types. Only the OTUs with at least 1% relative abundance were included for statistical analysis based on a FDR-corrected Kruskal-Wallis test (*P* <0.05). Overall, bulk soil and root types showed divergent abundances of different taxa at the order level (Fig. S3). Despite the large number of highly abundant orders (Fig. S3), taxonomic information is available for only a fraction of them (7 OTUs) (Fig. 2b). Root types tended to enrich the lowly abundant taxa from the soil. Taxa such as Pleosporales were widely enriched in all root types, and Agaricales were significantly enriched in lateral roots, whereas Chaetothyriales were significantly enriched in axial roots (Fig. 2b). Moreover, the taxonomic composition of free-living fungi in bulk soil was more complex than in roots (Fig. 2c). We further calculated the Shannon index as a measure of fungal diversity. At LP, the taxonomic diversity was higher in lateral roots than in axial roots, whereas, at HP the taxonomic diversity was similar for both root types (Fig. 2c). A dendrogram of differentially abundant phyla, normalized by *Z*-score across all data sets, suggested that taxonomic compositions were mainly separated by axial and lateral root types and only to a minor degree by P status (Fig. 2d). This indicated that root fungal community composition was stronger more strongly influenced by the host root types and less by P status. Still, under LP, the relative abundance of OTUs representing the phyla Zygomycota and Ascomycota was significantly higher in LR as compared to axial roots, while axial roots accumulated more OTUs for Chytridiomycota under this soil condition than lateral roots (Fig. 2e). Under HP, lateral roots showed a significantly higher abundance of Basidiomycota and Glomeromycota as compared to axial roots, whereas under HP conditions none of the large fungal phyla was enriched in axial roots (Fig. 2f).

### Maize root types are differentially colonized by AMF under field conditions

To monitor in detail whether maize root types grown in the field differ in their colonization levels by AM fungi, we microscopically quantified colonization frequency of axial roots and 1^st^ and 2^nd^ order lateral roots (Figs. S4, S5). This revealed that the 1^st^ order lateral roots were more strongly colonized than axial roots or the 2^nd^ order lateral roots under both P conditions (Fig. 3a). However, under LP conditions 1^st^ order lateral roots and axial roots were significantly more colonized than under HP condition, while surprisingly the P level did not significantly affect colonization of 2^nd^ order lateral roots (Fig. 3a). To support the notion that mycorrhizal colonization might be linked to functional differences among root types, the genes encoding Pi transporters of the PHT1 family including *ZmPht1;2*, *ZmPht1;4*, *ZmPht1;5*, *ZmPht1;6*, *ZmPht1;9*, *ZmPht1;10*, *ZmPht1;11* and *ZmPht1;13* were identified in the RNA-Seq dataset. Transcript accumulation of all *Pht1* genes was negatively correlated with P availability (Fig. 3b). All genes were preferentially expressed in lateral roots than in axial roots reflecting the stronger involvement of lateral roots in P uptake and AM symbiosis. Furthermore, axial roots exhibited larger variations in genes expression in response to external P changes than lateral roots (Fig. 3b). The differential colonization of root types at different P levels was confirmed by transcript accumulation of the maize AM marker gene *AM3* (Fig. 3b). Taken together, maize roots types displayed divergent AM colonization frequencies, which depended on external P status. However, independent of the P status 1^st^ order lateral roots were preferentially colonized.

**Fig. 3.**
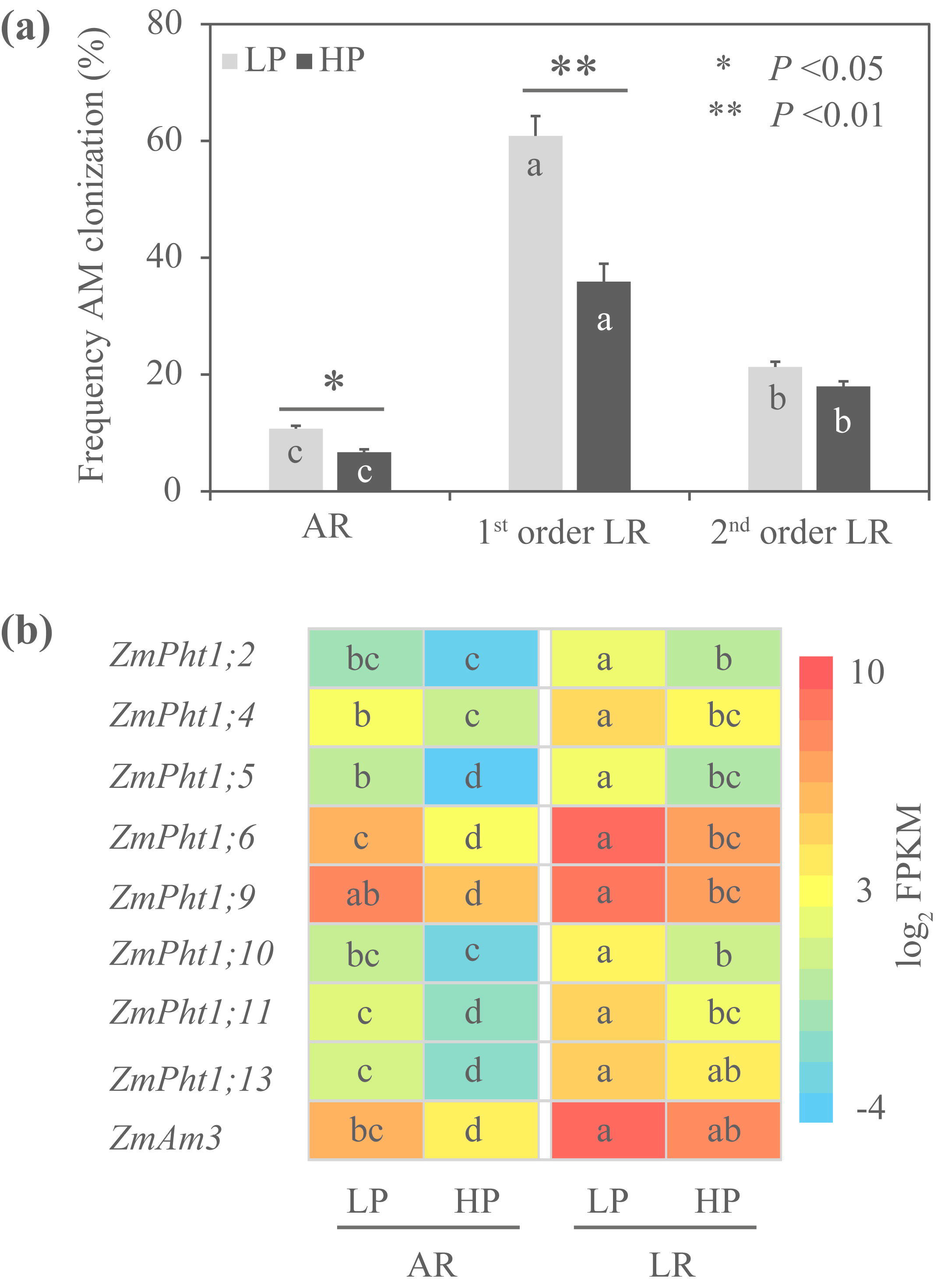
Colonization by AMF and expression of *ZmPht* genes in different root types of maize grown under LP and HP conditions. (a) Frequency of AM colonization (%) under HP (high phosphate) and LP (low phosphate) conditions. Asterisks denote significant differences between low and high P levels for a given root type according to paired Student’s *t* tests (\**P* <0.05; \*\**P* <0.01). Different letters above each bar indicate significantly differences among all conditions as assessed by post-hoc test (*P* <0.05). (b) Accumulation of *ZmPht* transcripts. The expression values were normalized by log_2_ transformed fragments per kilobase of transcript per million reads (FPKM). Significant differences are indicated by different letters (Tukey’s post-hoc test, *P* <0.05) and were calculated independently for each gene.

### Divergent transcriptomic responses of maize root types to soil P-availability in the field

The transcriptomes of axial and lateral roots were analyzed for significant differences at each P condition using a log-transformed linear model. This survey revealed that differential gene expression between axial and lateral roots was partially dependent on the P condition (Fig. 4a). In total, 6,955 genes were differentially expressed between axial and lateral roots (Fig. 4a). Among those, 2,724 transcripts accumulated differentially for both P treatments whereas 954 (14%) differed specifically at LP conditions, while a larger number of 3,277 (47%) transcripts differed specifically at HP conditions (Fig. 4a). The complete list of differentially expressed genes is provided in Dataset S5. Based on previously characterized phosphate starvation responsive (PSR) genes of maize (He *et al*., 2015), 25 candidate PSR genes significantly responded to phosphate levels (Fig. 4b), while the other genes determined by He **et al*.,* (2015) were not expressed or did not differ in expression between the different conditions. These candidate PSR genes clustered into two groups. Group I includes 17 genes induced by low phosphate in both root types. Group II includes eight genes, which also respond to low phosphate but are in addition significantly higher expressed in lateral roots as compared to axial roots (Fig. 4b).

**Fig. 4.**
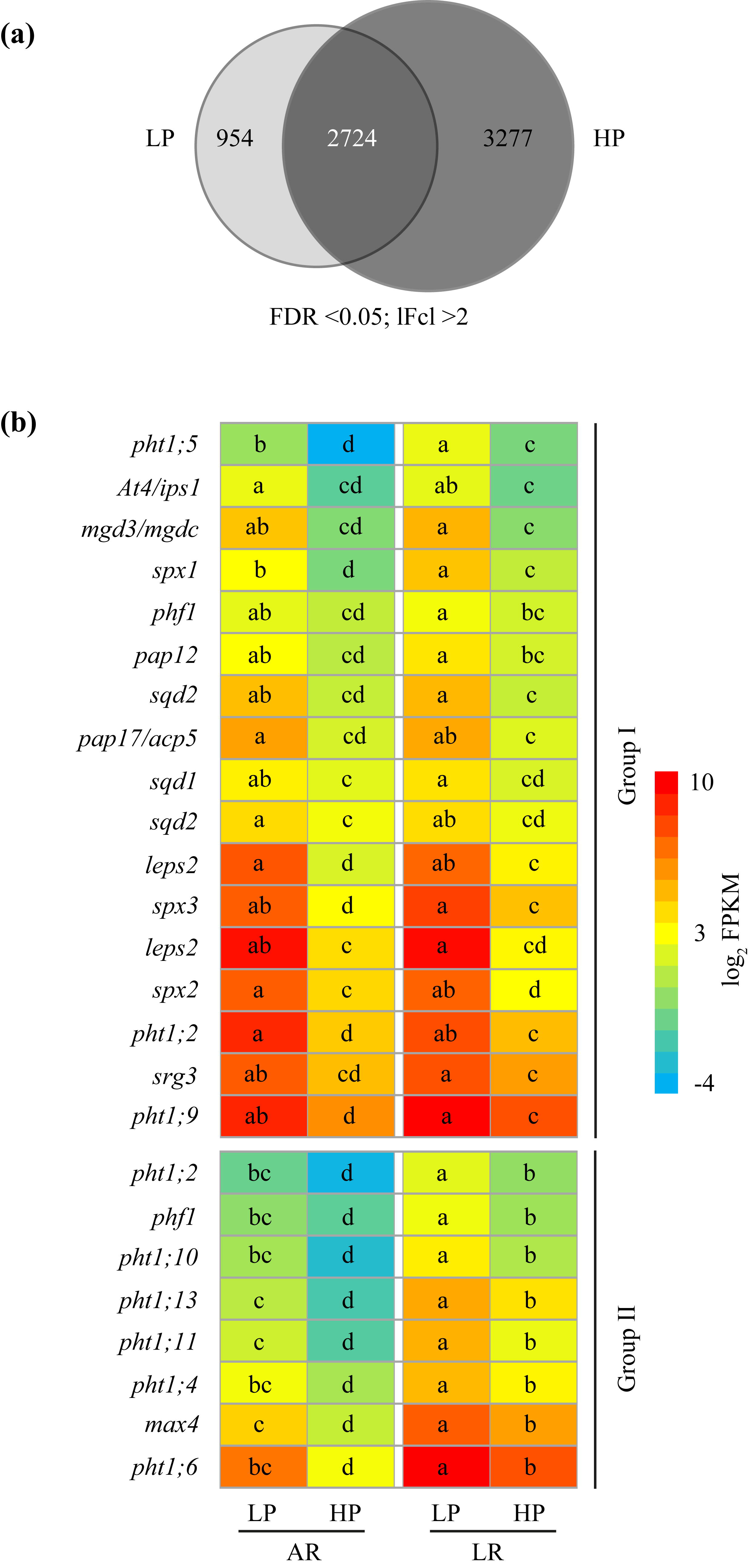
Root type-specific transcriptional responses to phosphate status. (a) Venn diagram illustrating the number of genes, which are differentially expressed between root types under low or high P conditions. FDR <0.05; |Fc| >2. (b) Expression pattern of phosphate starvation response (PSR) genes in maize root types. In total, 25 PSR genes (He *et al*., 2015) were detected from the list of genes, which are differentially expressed among root types. The expression values were normalized by log_2_ transformed FPKM. Significant differences are indicated by different letters (Tukey’s post-hoc test, *P* <0.05) and were calculated independently for each gene.

Genes differentially expressed between lateral and axial roots under specific P-levels were assigned to MapMan functional categories to compare the distribution of over-and under-represented functional classes between root types under low and high P (Tables S5-9). Based on Fisher’s exact test, the functional category “signalling” was exclusively enriched at low P (Fig. 5a). In contrast, the pathways “metal handling” and “DNA” were only enriched under high P conditions (Fig. 5a). The genes which were only differently expressed between the root types at LP (954 genes) and at HP (3,277 genes; Fig. 4a) were assigned to two classes reflecting the root type in which they were higher expressed (Fig. 5b). For the four resulting groups of genes, enrichment of MapMan functional categories was calculated. At LP, the bins “cell wall” and “signalling” were significantly overrepresented in axial roots (Fig. 5b). At HP the MapMan bins “cell wall", “secondary metabolism” and “stress” were significantly overrepresented in lateral roots (Fig. 5b). For example, twelve genes (GRMZM2G015654, GRMZM2G096268, GRMZM2G103128, GRMZM2G127184, GRMZM2G333274, GRMZM2G387087, GRMZM2G470010, GRMZM2G471594, GRMZM2G857459, GRMZM5G870571, GRMZM5G875445, GRMZM5G858456) related to hemicelluloses synthesis and enriched in “cell wall” category were exclusively upregulated in lateral roots (Table S8). Moreover, a number of defense-related, disease resistance and pathogenesis-related genes were enriched in the MapMan bin “stress” and were overrepresented in lateral roots (Table S9). Interestingly, at LP the category “cell wall” was overrepresented in axial roots while at HP the same category was overrepresented in lateral roots (Fig. 5b).

**Fig. 5.**
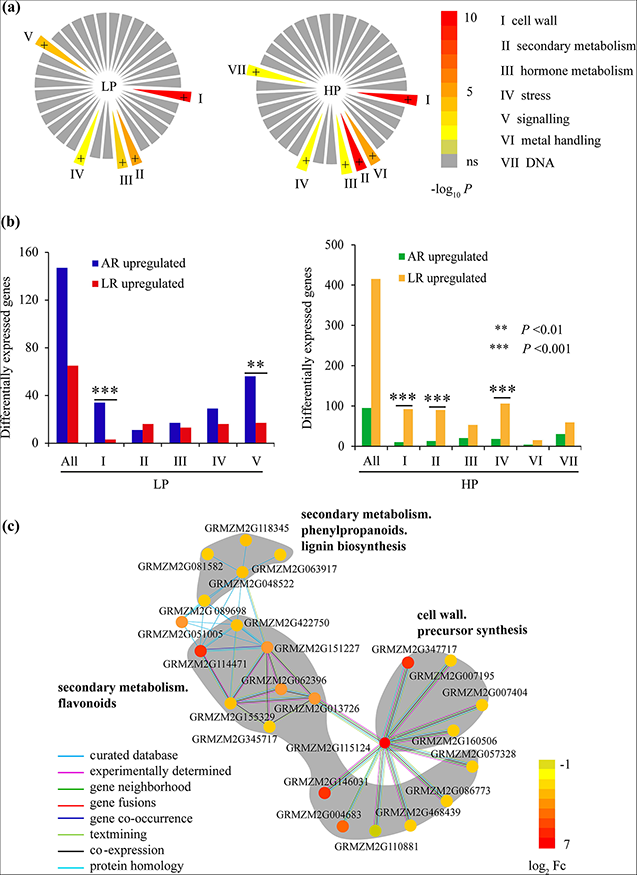
Enrichment of functional categories in maize root transcriptomes under LP and HP conditions. (a) Significantly enriched MapMan pathways under low and high P conditions detected by Fisher’s exact test (*P* <0.05). (b) Functional categories in which the ratio of preferentially expressed genes differed significantly from the ratio of all expressed genes from the enriched MapMan categories. The expected number of genes for each functional category was calculated based on the distribution of functional categories among all expressed genes. To determine, which genes were significantly overrepresented in which individual MapMan category, a χ^2^ test for independence with Yates' continuity correction was performed. \*\**P* <0.01; \*\*\**P* <0.001. (c) Network view of genes with high confidence scores of >0.7 generated using the STRING v10 prediction algorithm (Szklarczyk *et al*., 2015). Color codes from gray to red indicate expression levels of transcripts enriched in axial roots normalized as log_2_ logarithm of fold changes (log_2_Fc) under high phosphate conditions.

To uncover links between the genes belonging to given functional categories, we constructed a functionally connected network with the aid of the STRING algorithm at a high confidence level of >0.7. Under LP conditions transcripts (GRMZM2G017678, GRMZM2G031311, GRMZM2G042179, GRMZM2G052357, GRMZM2G063949, GRMZM2G161233, GRMZM2G429118) encoding previously characterized interacting proteins were strikingly enriched in axial roots and corresponded to cell wall biosynthesis and metabolism (Table S5). In contrast, under HP conditions eleven secondary metabolism pathways were enriched (Tables S8-10), of which the three pathways “cell wall precursor synthesis", “secondary metabolism flavonoids” and “secondary metabolism phenylpropanoids lignin biosynthesis” were biologically connected (Fig. 5c; Table S11).

## Discussion

Different root types of maize display distinct molecular (Gutjahr *et al*., 2015; Yu *et al*., 2016b) and physiological characteristics (Tai *et al*., 2016). Here we tested the hypothesis that axial and lateral roots of maize provide differentiated niches for root inhabiting microbes by root-type specific profiling fungal communities in field grown plants and the surrounding soil. In parallel, we examined the transcriptomes of these root types, to demonstrate root type-specific differences in physiology, metabolism and signalling under field conditions.

We found differences in the fungal communities between root systems and bulk-soil, confirming that roots provide a selective environment for microbes (Bulgarelli *et al*., 2012; de Souza *et al*., 2016). Strikingly, more than half of the OTUs identified inside maize roots were specific for one or the other root type (Fig. 2a). Furthermore, the relative amount of OTUs belonging to certain fungal phyla, the Agaricales, Sordariales, Ascomycota sp., Chaetothyriales, Mortierellales and Pleosporales, was significantly different between the two maize root types (Fig. 2b). This demonstrates that maize root types influence the endophytic fungal community in a realistic field situation. Differences in bacterial and fungal communities between roots and leaves and/or stalks have previously been described for *Arabidopsis* and sugar cane (Bai *et al*., 2015; de Souza *et al*., 2016). We show that fungal communities diverge even among different parts of the same plant organ (the root system).

Phosphate nutritional status of the plant or the ability of the plant to perceive phosphate have been shown to strongly affect the composition of the bacterial root microbiome (Castrillo *et al*., 2017) and the ability of a fungal endophyte to colonize Arabidopsis (Hacquard *et al*., 2016; Hiruma *et al*., 2016). Here we found, that the soil phosphate level had a profound and root type-specific effect on the structure of fungal communities (Fig. 2a,b), suggesting that the phosphate status effects niches for fungal colonization in a root type-specific manner. In field-grown maize roots, phosphate influenced the ß-diversity of root inhabiting fungi specifically in LRs, which displayed higher ß-diversity at low P than high P, while in AR the ß-diversity was similar in both P conditions. Furthermore, the analysis of species composition suggests that LRs support a higher diversity of Ascomycota at LP and of Basidiomycota and Glomeromycota at HP, while ARs support a high diversity of Chytridiomycota at LP (Fig. 2d). A slight shift in the fungal community composition in response to differences in soil phosphate level was also observed in bulk soil, suggesting that some fungi are influenced by phosphate directly or indirectly for example through plant phosphate-status dependent root exudates (Yoneyama *et al*., 2013; Ziegler *et al*., 2016).

It has been demonstrated that the expanded capacity of AM roots to gain soil P by long-distance transport presents a major contribution to nutrient uptake in crops (reviewed in Sawers *et al*., 2008; Smith & Smith, 2011; Gutjahr & Paszkowski, 2013). We found a root type-specific distribution of AM colonization, with higher colonization levels in LRs as compared to ARs, as previously reported for rice (Gutjahr *et al*., 2015). The accumulation of AM related transcripts such as *Pht1;2, Pht1;5*, and *Pht1;6* correlated with colonization of lateral roots under different P regimes (Fig. 4e; Liu *et al*., 2016; Sawers *et al*., 2017). Increased AM colonization and induction of AM-specific Pi transporters genes *ZmPht1;5* and *ZmPht1;6* in lateral roots likely enhances P acquisition via mycorrhizal pathway under LP conditions (Willmann *et al*., 2013; Deng *et al*., 2014; Sawers *et al*., 2017).

High phosphate suppressed the amount of root colonization by AMF and the expression of AM marker and phosphate transporter genes, as previously reported (Fig. 3d-e, reviewed in Carbonnel & Gutjahr, 2014). Surprisingly, this condition leads to an increase in the diversity of AMF species, specifically in lateral roots. Root colonization by AMF is controlled by the plant and is suppressed if the fungus does not deliver phosphate (Javot *et al*., 2007). Based on this finding it was hypothesized that under low phosphate, plants promote root colonization by AMF species, which are most efficient in phosphate delivery and suppress less efficient species (Gutjahr & Parniske, 2017). In turn, it is possible that at higher phosphate levels, fungal species, which are less phosphate-efficient are permitted to colonize, possibly because they provide other services to the plant such as an increased biotic or abiotic stress resistance (Gianinazzi *et al*., 2010).

Simultaneous with fungal OTUs in maize root types, we determined more divergent transcriptomic differences among root types than among P treatments indicating that root type identity dominated the transcriptional profile (Fig. 1c). However, differential accumulation of transcripts related to phosphate starvation confirmed that the plants had responded to the phosphate treatment; and some of these transcripts accumulated differentially between axial and lateral roots (Fig. 4b). A number of transcripts were enriched in a root type-specific manner independent on the phosphate status, and root type specificity of the transcriptome was higher under HP than under LP (Fig. 4a). Under LP the functional categories “cell wall” and “signalling” were overrepresented in axial roots (Fig. 5a, b; Tables S5, 6). Enrichment of cell wall related transcripts in crown roots in comparison to lateral roots has also previously been observed in rice grown under controlled phytochamber conditions (Gutjahr *et al*., 2015). We demonstrate here that this also occurs in the field and in a second grass species, indicating that this is likely a general phenomenon. The increased accumulation of transcripts encoding cell wall precursors likely promotes a strengthened cell wall structure, which may explain the lower colonization of axial roots by AM fungi as compared to lateral roots under LP conditions (Fig. 4a-d; Table S5). Downregulation of transcripts related to cell wall modification associated with the establishment of AMF symbioses has also been observed in rice (Gutjahr *et al*., 2015; Fig. 4a-d; Table S5), consistent with the higher AMF colonization (Figs 3a). At HP we observed an enrichment of transcripts belonging to the MapMan functional categories “cell wall", “secondary metabolism” and “stress” in lateral roots (Tables S7-9). For example, twelve genes associated with hemicelluloses synthesis were exclusively upregulated in lateral roots (Table S8). The most important biological function of hemicelluloses is their contribution to strengthening the cell wall by tethering cellulose microfibrils (Scheller & Ulvskov, 2010). Interestingly, a maize *Pht1;6* knock-out, which is perturbed in mycorrhizal phosphate uptake showed lower expression of cell wall related genes than the wild-type (Willmann *et al*., 2013), confirming that also in phosphate poor soils higher P levels (as in the wild-type) support activation of cell wall processes.

In addition, the transcripts encoding proteins, which may be involved in the inhibition of fungal pathogens are enriched in lateral roots (Table S8). We observed strong inductions of defense related transcripts enriched in the MapMan category “stress” such as *nucleotide-binding, leucine-rich repeat (NB-LRR)* genes (GRMZM2G045027, GRMZM2G116335, GRMZM2G156351, GRMZM2G440849, GRMZM5G880361), the *polygalacturonase-inhibiting proteins (PGIPs)* genes (GRMZM2G025105, GRMZM2G099295, GRMZM2G129493) and the *Respiratory burst oxidase homolog* (*Rboh*) gene family (GRMZM2G037993, GRMZM2G065144, GRMZM2G300965, GRMZM2G358619), which have all been implicated in defense against fungal pathogens (Ferrari *et al*., 2003; Torres & Dangl, 2005; Gao *et al*., 2011; Table S9). Furthermore, lignin biosynthesis genes such as *4CL1* (*4-coumarate: CoA ligase 1*; GRMZM2G048522, GRMZM2G174732) (Fraser & Chapple, 2011), the expression of which has been shown to correlate with lignin deposition, may be involved in protecting lateral root initiation sites against pathogen infiltration (Fig. 5c; Tables S10, 11). Taken together, these results indicate that distinct biological processes are enriched in divergent root types at different external P concentrations, due to a specific difference in the adaptive responses among the root types to external P levels (Figs 2c, d, 5; Tables S7, 8).

In Arabidopsis, phosphate sufficiency was associated with increased defense gene expression in roots. This was made responsible for changes in a synthetic bacterial rhizosphere and root community (Castrillo *et al*., 2017) and suppression of the endophytic fungus *Colletotrichum tofieldiae*, which delivers phosphate under LP conditions (Hacquard *et al*., 2016; Hiruma *et al*., 2016). We hypothesize that phosphate-induced defense responses in lateral roots could also be responsible for the fungal community shifts we observed in field-grown maize roots grown at HP (Fig. 5). We consider it likely that specific reprogramming of root types in response to the soil phosphate condition determines the root type-specific fungal community composition (see summary in Fig. 6). However, we do not exclude that part of the observed transcript accumulation occurs in response to the colonization by the fungi. In addition it is most likely that also the community composition of other microbes such as bacteria, is influenced by root type specific niches and phosphate conditions and that in turn these communities have an impact on the root transcriptome and physiology. It remains to be determined to which extent transcriptome shifts in response to environmental stimuli such as phosphate are causative for fungal colonization or in turn influenced by the inhabiting fungi. Nevertheless, the data presented here provide a framework for novel research aiming at an understanding of root type-specific responses to biotic and abiotic factors and will guide future efforts to improve plant growth and fitness, through application of soil microbes.

**Fig. 6.**
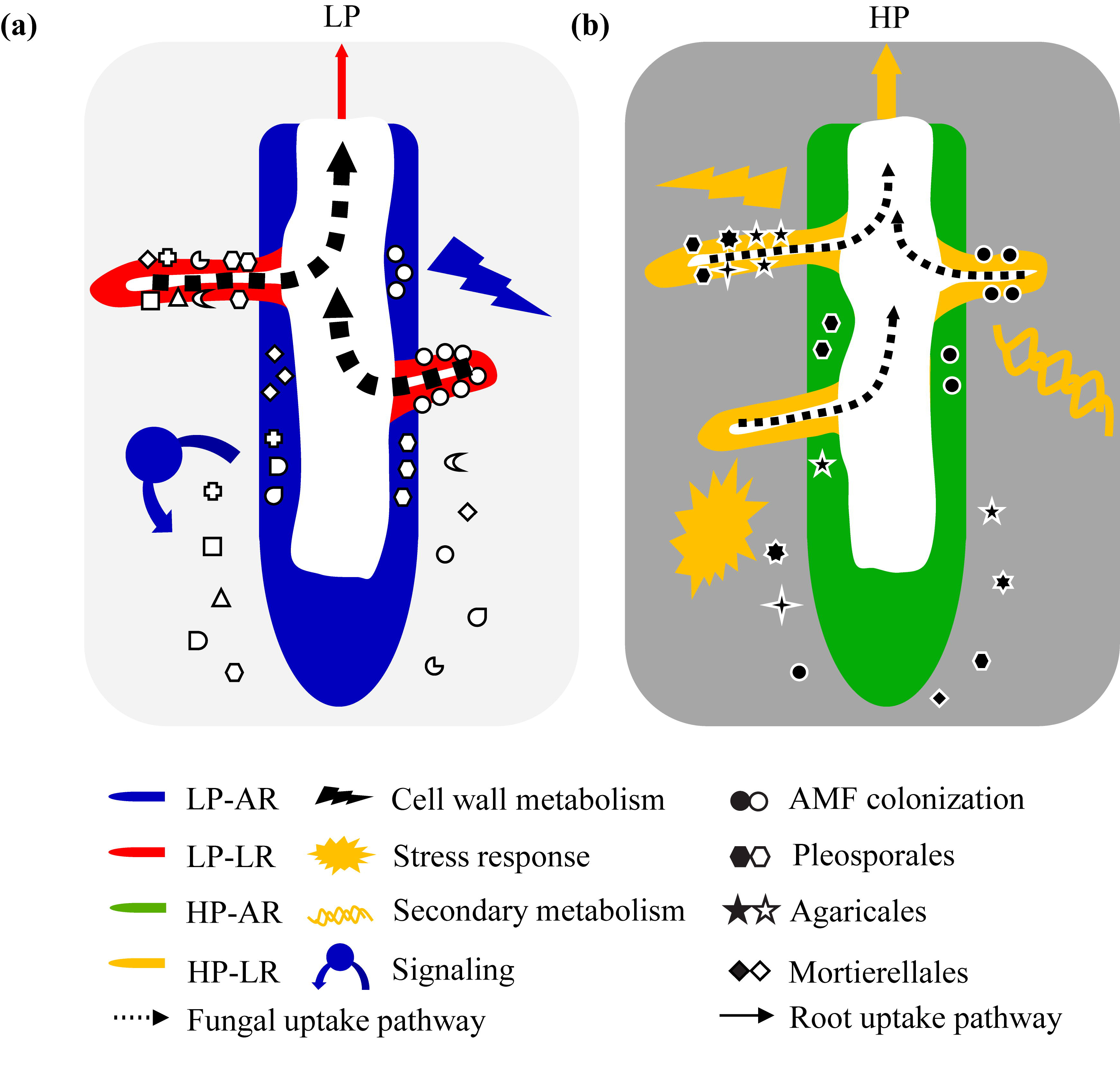
Summary of phosphate-dependent enrichment of functional transcriptome categories and of host-inhibiting fungal communities in field-grown root types. Under P-deficient conditions, selective colonization by nutrient-delivering AMF and higher expression of P foraging genes in lateral roots might contribute to P acquisition of the shoot (Figs 3, 4b). Enrichment of cell wall metabolism in axial roots might explain the relatively higher colonization in axial roots under LP conditions compared to HP conditions (Fig. 5). Under P-sufficient conditions, P acquisition is mainly contributed by the host roots themselves with less fungal colonization (Fig. 3). In contrast, enrichment of defense-related biological processes in lateral roots might be induced by high P accumulation in both, bulk soil and plant shoot (Fig. 5; Tables S1, S2, S4). Solid arrows of different thickness indicate the degrees of P acquisition by direct phosphate uptake. Dotted arrows indicate the contribution of P acquisition via AM. Fungal communities are represented by diverse shapes of different symbols, of which the black and white “dots” indicate AMF. AR, axial roots; HP, high phosphate; LP, low phosphate; LR, lateral roots.

## Acknowledgements

Root research in C.L.’s laboratory is supported by grants of the State Key Basic Research and Development Plan of China (No. 2013CB127402) and the Innovative Group Grant of the National Natural Science Foundation of China (NSFC) (No. 31421092). Maize root research in F.H.’s laboratory is supported by the Deutsche Forschungsgemeinschaft (DFG) and the Federal Ministry of Education and Research (BMBF). C.G. is supported by the Emmy Noether program (GU1423/1-1) of the DFG and her maize field research by BayKlimaFit of the Bavarian Ministry of Environment and Consumer Protection.

## Author Contribution

C.L and F.H. conceived and designed this field experiment. C.W. collected the field samples and performed the arbuscular mycorrhizal staining experiments. P.Y. analyzed the fungal ITS data. P.Y., J.A.B. and H.T. analyzed the transcriptome data. C.G and F.H. contributed to data interpretation. P.Y. and C.G. wrote the manuscript.

## Supporting Information

**Fig. S1. RNA quality assessment of 16 maize root type samples collected from the field.** AR, axial root; HP, high phosphate; LP, low phosphate; LR, lateral root; RIN, RNA integrity number.

**Fig. S2. Diversity index (alpha rarefaction) of ITS sequences.** Chao 1 is estimated as the species abundance. AR, axial root; BS, bulk soil; HP, high phosphate; LP, low phosphate; LR, lateral root.

**Fig. S3. Relative abundance of fungal taxa in maize root types at two phosphate levels.** Orders with an average relative abundance lower than 1% were attributed to “others". Statistical significance was tested using the Kruskal-Wallis test (FDR-corrected *P* <0.05) at the order level. Un: Unknown. The same taxa names indicate different OTU identities. AR, axial root; BS, bulk soil; HP, high phosphate; LP, low phosphate; LR, lateral root.

**Fig. S4. Distribution of AM colonization in maize root types along the longitudinal root axis.** Axial and lateral root types were dissected along the whole axial root into 5 cm pieces and stained by Trypan blue under LP conditions (a) and HP conditions (b). Dotted lines across the axial roots indicate the positions at which the root segments were collected and stained. Each pair of pictures above the dotted lines are representative for lateral and axial root types from different segments of the whole axis. The red line indicates lateral roots grown under LP conditions. The blue line indicates axial roots grown under HP conditions. The yellow line indicates lateral roots grown under HP conditions. The green line indicates axial roots grown under HP conditions. Asterisks denote significant colonization differences between axial and lateral root types given a specific P level according to paired Student’s *t* tests (\**P* <0.05; \*\**P* <0.01). AM, arbuscular mycorrhiza; HP, high phosphate; LP, low phosphate.

**Fig. S5. Visualization of fungal structures in axial root, 1^st^ and 2^nd^ order lateral roots stained with Trypan blue.** AR, axial root; 1^st^LR, 1^st^ order lateral roots; 2^nd^ LR, 2^nd^ order lateral roots; HP, high phosphate; LP, low phosphate. Scale bar = 200 µm.

**Dataset S1. Assembly results and quality control of fungal ITS sequencing.**

**Dataset S2. Overview of RNA-Seq output, mapping results and alignments to the B73 reference genome.**

**Dataset S3. Complete list of 27,375 expressed genes.**

**Dataset S4. Complete list of relative abundance of fungal enriched OTUs and taxonomy information.**

**Dataset S5. Complete list of 6,955 differentially expressed genes under both LP and HP conditions.** HP, high phosphate; LP, low phosphate. The differentially expressed were divided into three groups according to the Fig. 4a. Light grey color indicates the root type-specific differentially expressed genes under the LP conditions. Dark grey indicates the root type-specific differentially expressed genes under the HP conditions.

